# Resolving early cochlear inflammation prevents lasting damage from noise exposure

**DOI:** 10.64898/2026.07.11.737965

**Authors:** Lauren Barbush, Klim Fedorchuk, Iman Ezzat, Vijayprakash Namakkal Manickam, Dinesh Gawande, Ayden Chavez, Marisa Zallocchi

## Abstract

Noise-induced hearing loss (NIHL) is a leading cause of permanent hearing impairment worldwide, yet no pharmacological therapies are currently available to prevent or treat this disorder. Although inflammation is increasingly recognized as a key contributor to cochlear degeneration, the therapeutic potential of targeting early inflammatory signaling remains poorly understood. Here, we combined phenotypic screening in zebrafish with mechanistic and functional validation in complementary mouse models to identify quinoxaline derivatives with otoprotective activity following acoustic trauma. Lead compounds preserved cochlear synapses and auditory function after moderate noise exposure, while one derivative also protected sensory hair cells in a model of permanent hearing loss. Mechanistic analyses demonstrated that this protection was associated with attenuation of early NF-κB signaling and modulation of the cochlear inflammatory response toward a reparative state, consistent with suppression of pathogenic innate immune activation before irreversible tissue damage occurred. Together, these findings identify early NF-κB-dependent inflammatory signaling as a therapeutically actionable mechanism in NIHL and establish quinoxaline derivatives as promising candidates for pharmacological intervention. More broadly, this work demonstrates the utility of a cross-species discovery platform for identifying therapies that preserve sensory function by targeting early inflammatory pathways.

## INTRODUCTION

More than 300 million people worldwide suffer from noise-induced hearing loss (NIHL)^1^. Members of the military are particularly vulnerable due to blast exposures and acoustic trauma from artillery fire or explosives, which can produce extreme sound pressures^1–3^. The effects of intense noise are rapid, with phenotypic abnormalities evident immediately after exposure^4–11^. Severe acoustic trauma is associated with cochlear hair cell (HC) degeneration and neuronal excitotoxicity driven by excessive accumulation of the neurotransmitter glutamate^1 5 8 11–13^. Overactivation of AMPA receptors by glutamate leads to swelling and bursting of postsynaptic nerve fibers, ultimately contributing to HC loss^14–17^. Furthermore, studies in animal models have shown that noise-induced synaptic injury precedes HC death and neuronal degeneration^4 5 8 11–13 17^, an observation that may explain the speech perception difficulties in noisy environments experienced by some individuals even while they retain normal hearing thresholds^11 15^. Despite the considerable global impact of NIHL, its molecular mechanisms remain poorly understood, and no effective therapeutic intervention currently exists to prevent or alleviate this condition.

Prior research from our laboratory supports the therapeutic potential of quinoxaline (Qx) derivatives for treating ototoxin-induced damage and possibly, NIHL^18 19^. Unlike agents currently in pre-clinical or clinical trials, which act at later stages during reactive oxygen species (ROS) generation, after irreversible oxidative damage to the auditory system may have already occurred^20^, quinoxaline and some of its derivatives act upstream by blocking activation of the nuclear factor κB (NF-κB) pathway^21^. Upregulation of NF-κB has been observed in the mouse inner ear after acoustic trauma^22–26^; therefore, by preventing NF-κB activation, quinoxaline derivatives may allow cells to recover from ototoxic insult before irreversible damage takes place. Interest in this chemical class has grown due to its broad biological activity and therapeutic potential, including anti-cancer, anti-microbial, and neuroprotective properties^16 27–30^. Quinoxaline derivatives also possess several properties that make them attractive candidates for drug discovery: structural malleability for medicinal chemistry, the ability of many derivatives to cross the blood-labyrinth barrier (BLB)^27^, and water solubility and stability at physiological pH^16 18 21^. Moreover, several quinoxaline derivatives are already FDA-approved for use in the pharmaceutical and food industries^31^, which could facilitate their repurposing for auditory therapeutics. This approach is associated with shorter development timelines, reduced costs, and higher approval rates than *de novo* drug discovery^32^.

Here, we screened a library of Qx derivatives using a pharmacologically induced zebrafish model that mimics acoustic trauma^5^. Zebrafish HCs and their associated supporting cells and neurons are morphologically and functionally analogous to their mammalian counterparts and exhibit similar responses to ototoxic injury^33–36^. Thus, lead compounds showing protective effects in zebrafish could be advanced for testing in mouse models of ototoxic trauma. Through this pipeline, we identified Qx34 and Qx62 as our top lead candidates, conferring protection against noise-induced synaptopathy and HC death. Moreover, since Qx62’s mechanism of action involves, at least in part, attenuation of the inflammatory response through the modulation of NF-κB pathway, it may also hold therapeutic potential for other forms of acquired hearing loss. Thus, Qx62’s anti-inflammatory properties could provide broader insights into diseases in which inflammation is a key pathogenic driver.

## RESULTS

### Screening of quinoxaline derivatives in a zebrafish model of excitotoxic damage

To identify compounds with the potential to protect against excitotoxicity, we screened a previously published library of Qx derivatives in zebrafish (**Supp. Fig. 1**)^19^. Wild type larvae (5-6 days post-fertilization - dpf) were exposed to 600µM kainic acid (KA) for 1 hour, followed by a 2-hour incubation with one of the Qx derivatives. After fixation and immunostaining with the HC marker otoferlin, neuromast HCs were manually counted and compared with those in animals treated with kainic acid alone. A total of 69 derivatives (**Supp. Fig. 1**), including the original quinoxaline molecule (Qx1), were tested; of these, 25 derivatives (∼36%) showed significant protection at two or more concentrations (**Supp. Fig. 2** and **Fig. 1**).

**Figure 1.**
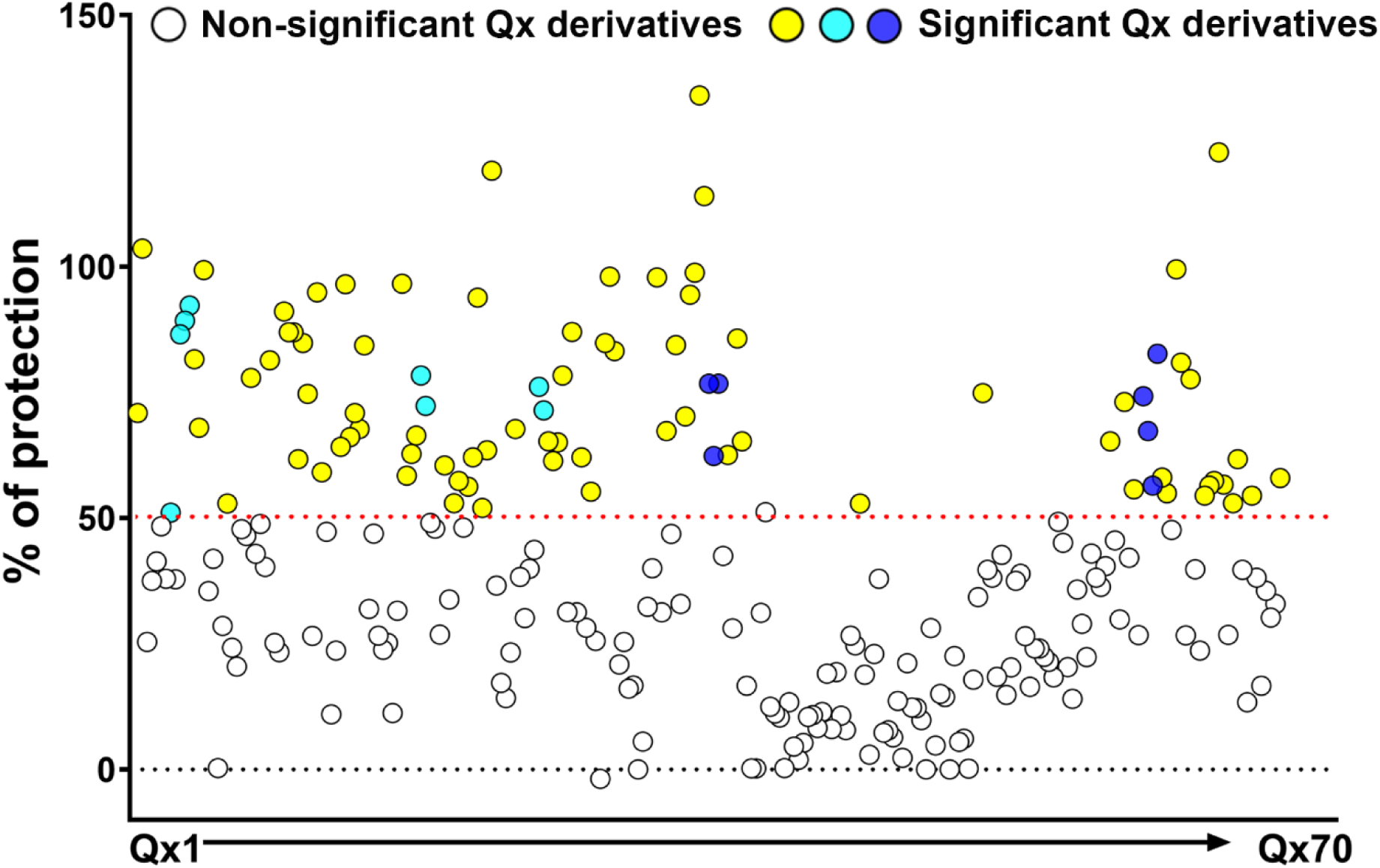
Screening of quinoxaline derivatives in a zebrafish model that mimics excitotoxicity. Five- to six-day-post-fertilization zebrafish were incubated with 600 µM kainic acid for 1 hour, followed by a 2-hour incubation with one of the quinoxaline derivatives (Qx1-Qx70) at concentrations ranging from 1 nM to 100 µM. Neuromast hair cells were immunostained for otoferlin and quantified under a fluorescence microscope. Results are expressed as the percentage of protection relative to the kainic acid-only group (0%), with the vehicle control (DMSO alone) defined as 100% protection. Each dot represents one zebrafish. White dots indicate Qx derivative/concentration combinations that did not provide significant protection compared to kainic acid-only group. Yellow, cyan and blue dots indicate combinations that provided significant protection. Cyan dots correspond to Qx3, Qx17 and Qx23. Blue dots correspond to Qx34 and Qx62. The neuromasts analyzed were MI1, MI2 and O2. Statistical analysis was performed using one-way ANOVA followed by Dunnett’s *post hoc* test. N=5-6 fish per group, with each fish representing one biological replicate.

Because synaptic overstimulation caused by glutamate release and accumulation leads to ribbon synapse loss^5 17^, we next assessed whether any of these 25 compounds protected the ribbon synapses. For this purpose, we used the *Tg(NeuroD-GFP)* zebrafish line^37^, which expresses GFP in the afferent neuronal fibers. Larvae were incubated with kainic acid, followed by a 30-minute incubation with one of the derivatives at its lowest protective concentration. Animals were fixed and immunostained for GFP and the presynaptic marker Ribeye B (**Fig. 2**). Quantification of pre-synaptic ribbons (**Fig. 2A**) revealed that only five derivatives (20%) - Qx3, Qx17, Qx23, Qx34, and Qx62 (**Fig. 2B-O**) - significantly protected against kainic acid-induced synapse loss. Notably, these compounds also reduced or eliminated the characteristic afferent terminal swelling associated with excitatory damage (**Fig. 2B-O**, dotted frames). These findings indicate that the protective effects of Qx derivatives extend to both pre- and postsynaptic components during excitotoxicity.

**Figure 2.**
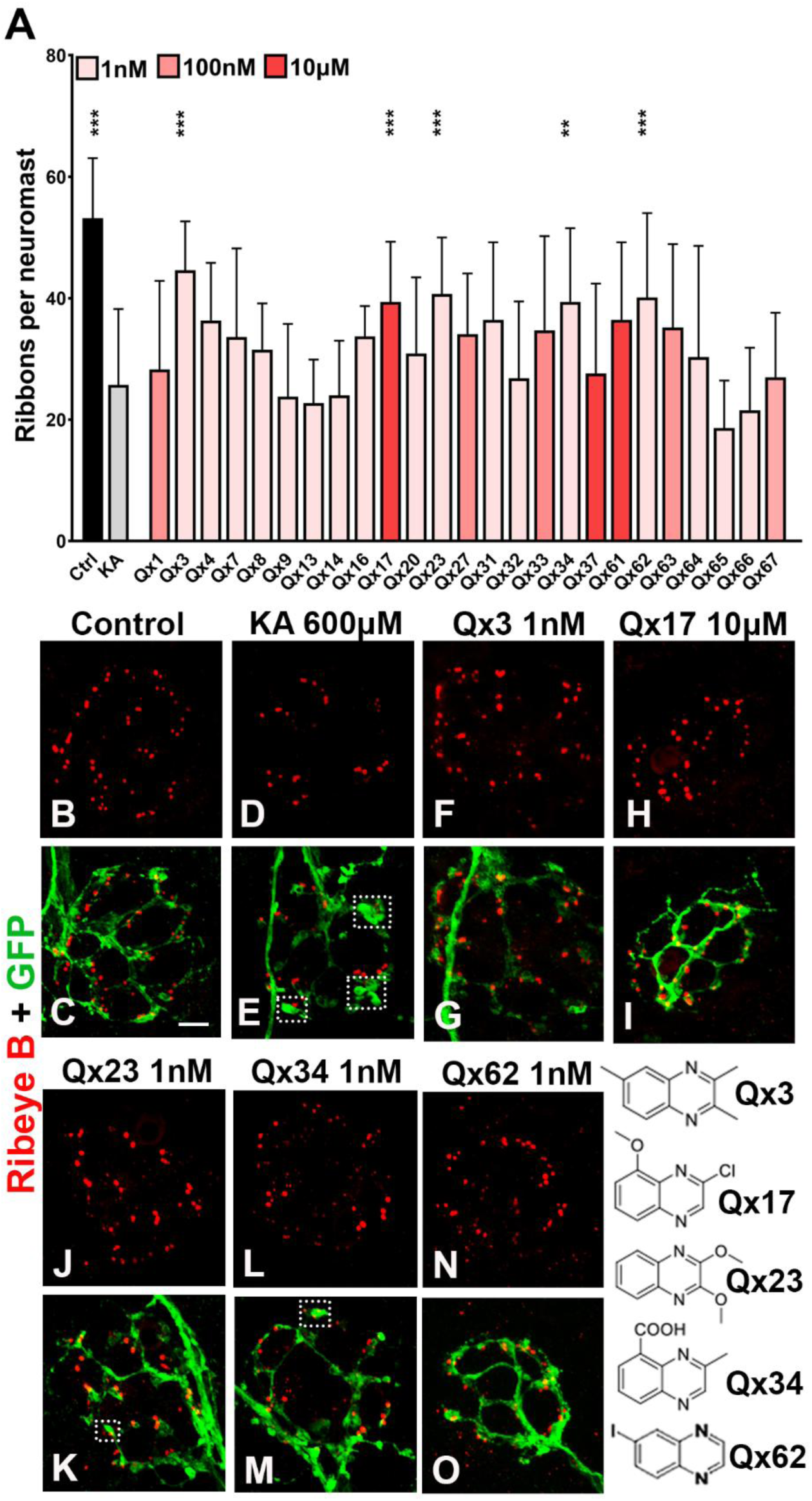
Quinoxaline derivatives protect against ribbon synapse loss. Five- to six-day-post-fertilization *Tg(NeuroD-GFP)* larvae were incubated with 600 µM kainic acid for 1 hour, followed by a 30-minute incubation with one of the quinoxaline derivatives at the lowest protective concentration identified in the screening. **A:** Results are expressed as the mean±SD of the number of pre-synaptic ribbons per neuromast. Black: vehicle control (Ctrl; DMSO). Grey: kainic acid-only (KA) group. Pink: KA + Qx derivative. **B-O:** Representative images of larvae immunostained for GFP (green) to label the afferent fibers and ribeye B (red) to identify the hair cell pre-synaptic ribbons. Each dot represents one zebrafish. Dotted rectangles indicate swelling of the neuronal terminals. Scale bar: 5 µm. The chemical structures of the Qx derivatives that protected the ribbon synapses are shown on the right. N=5-6 fish per group, with each fish representing one biological replicate. Statistical analysis was performed using one-way ANOVA followed by Dunnett’s *post hoc* test. **P<0.01, ***P<0.001 compared to KA alone.

In summary, our zebrafish screen identified five quinoxaline derivatives that protected against HC damage, ribbon synapse loss, and neuronal terminal swelling. These compounds were selected for further evaluation in a mouse model that more closely replicates noise-induced damage.

### Assessing the therapeutic potential of quinoxaline derivatives against noise-induced synaptopathy

We next tested these five derivatives in a mouse model of noise-induced synaptopathy. Auditory brainstem responses (ABRs) and distortion product otoacoustic emissions (DPOAEs) were measured in 7-8-week-old CBA/CaJ mice before treatment (baseline) and again at 1 (PNE1), 7 (PNE7), and 21 (PNE21) days post-noise exposure (**Supp. Figure 3A**). Mice were exposed to 94dB SPL, 8-16 kHz noise for 2 hours and immediately injected intraperitoneally with a single dose of one of the Qx derivatives (25mg/kg b.w., dose determined for quinoxaline in unpublished pilot studies). Noise-exposed animals injected with vehicle (corn oil), as well as non-noise-exposed controls, were also included.

As expected, ABR and DPOAE thresholds were significantly elevated at PNE1 in all noise-exposed groups (**Fig. 3** and **Supp. Fig. 4**). Mice treated with Qx34 (**Fig. 3D**) or Qx62 (**Fig. 3E**) showed accelerated recovery; by PNE7, thresholds had largely returned to normal across most frequencies, with only 32kHz and 45kHz still showing significant elevation. In contrast, mice treated with Qx3 (**Fig. 3A**) or Qx23 (**Fig. 3C**) recovered more slowly, with significant threshold elevations persisting across all frequencies at PNE7. Qx17 proved ototoxic; threshold elevations persisted at PNE21 in both noise-exposed and non-noise-exposed animals that received Qx17 (**Fig. 3B**), and we discontinued its investigation.

**Figure 3.**
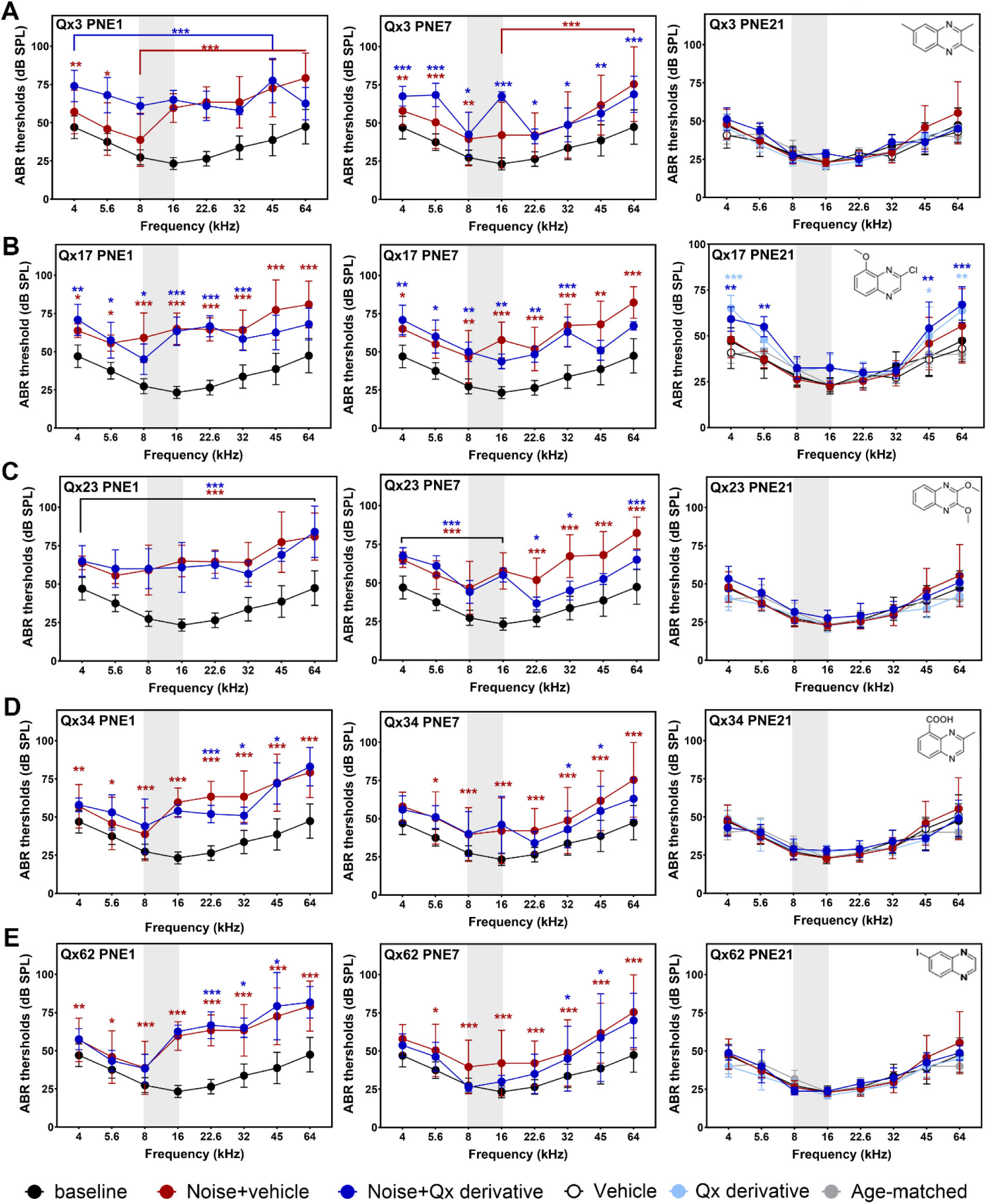
Qx derivatives accelerate ABR thresholds recovery after noise-induced synaptopathy. Seven- to eight-weeks old CBA/CaJ mice were exposed to 94dB SPL for 2 hours at 8-16 kHz and immediately IP injected with corn oil or one of the Qx derivatives at 25mg/kg b.w. ABR thresholds were measured at one (PNE1), seven (PNE7), and 21 (PNE21) days post-noise exposure. **A:** Qx3. **B:** Qx17, **C:** Qx23, **D:** Qx34 and **E:** Qx62. Results are expressed as mean±SD. Two-way ANOVA followed by Dunnett’s post-test for multiple comparisons. *P<0.05, **P<0.01, and ***P<0.001 Noise+vehicle (red asterisks) and Noise+Qx derivative (blue asterisks) *versus* baseline. ns = not significant. Number of animals: Noise+vehicle=11. Noise+Qx3=5. Noise+Qx17=6. Noise+Qx23=6. Noise+Qx34=5. Noise+Qx62=6. Vehicle=8. Qx derivative alone=6. Baseline=12. Age-matched=5. Gray shaded bar indicates the octave band noise.

Noise-induced synaptopathy is characterized by the recovery of ABR thresholds despite persistent loss of ribbon synapses, typically reflected in reduced wave-I amplitudes and increased latencies^9^. We therefore investigated whether Qx34 and Qx62 restored or preserved these parameters. At the 32kHz and 45kHz frequencies (**Fig. 4**), wave-I amplitudes (**Fig. 4A-D**) and latencies (**Fig. 4E-H**) in treated animals resembled those of non-noise-exposed controls, whereas noise-exposed, vehicle-treated mice showed significantly reduced amplitudes and prolonged latencies, confirming synaptic dysfunction. This was corroborated by immunostaining for the presynaptic ribbon marker CtBP2 (**Fig. 5**); at both the 32kHz (**Fig. 5A, 5C, 5E** and **5G**) and 45kHz (**Fig. 5B, 5D, 5F** and **5H**) frequency regions, noise-exposed animals showed a significant reduction in presynaptic puncta relative to vehicle-only controls, whereas mice treated with a single dose of Qx34 or Qx62 had puncta counts similar to vehicle-treated controls (**Fig. 5I**).

**Figure 4.**
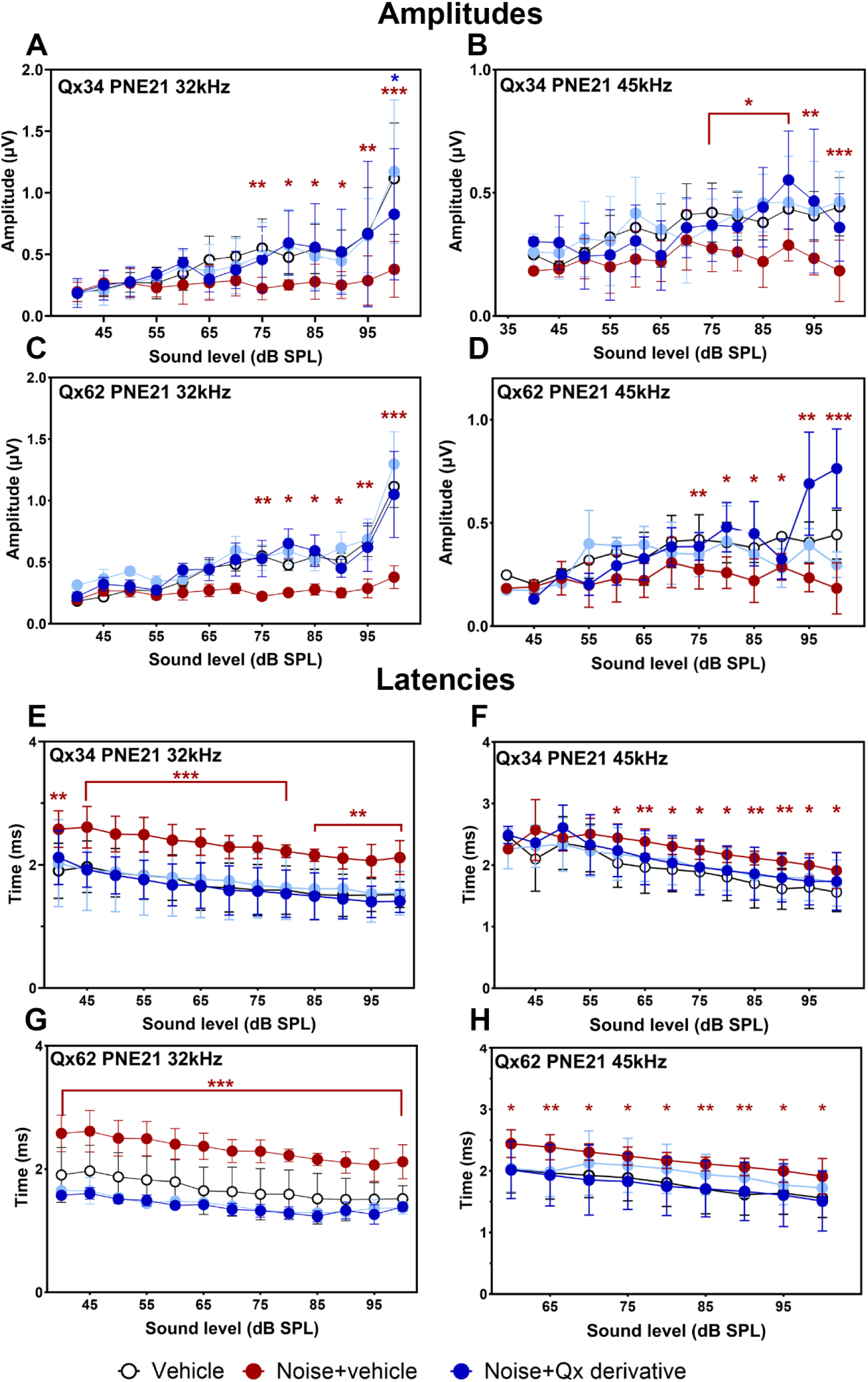
Qx34 and Qx62 preserve synaptic function after noise exposure. Wave-I amplitudes (**A-D**) and latencies (**E-H**) were measured 21 days after noise exposure in the Noise+vehicle (red), and Noise+Qx derivative (blue) groups at 32 kHz (**A, C, E, G**) and 45 kHz (**B, D, F, H**). White dots indicate vehicle-no noise group. Light blue dots indicated derivative-only group. Results are expressed as mean±SD. Statistical analysis was performed using two-way ANOVA followed by Dunnett’s multiple comparisons test, *P<0.05, **P<0.01, ***P<0.001. Red asterisks: Noise+vehicle *versus* Vehicle. Blue asterisks: Noise+Qx derivative *versus* Vehicle. Group sizes were as follows: Noise+vehicle, N=11; Noise+Qx34, N=5; Noise+Qx62, N=6; Vehicle, N=8.

**Figure 5.**
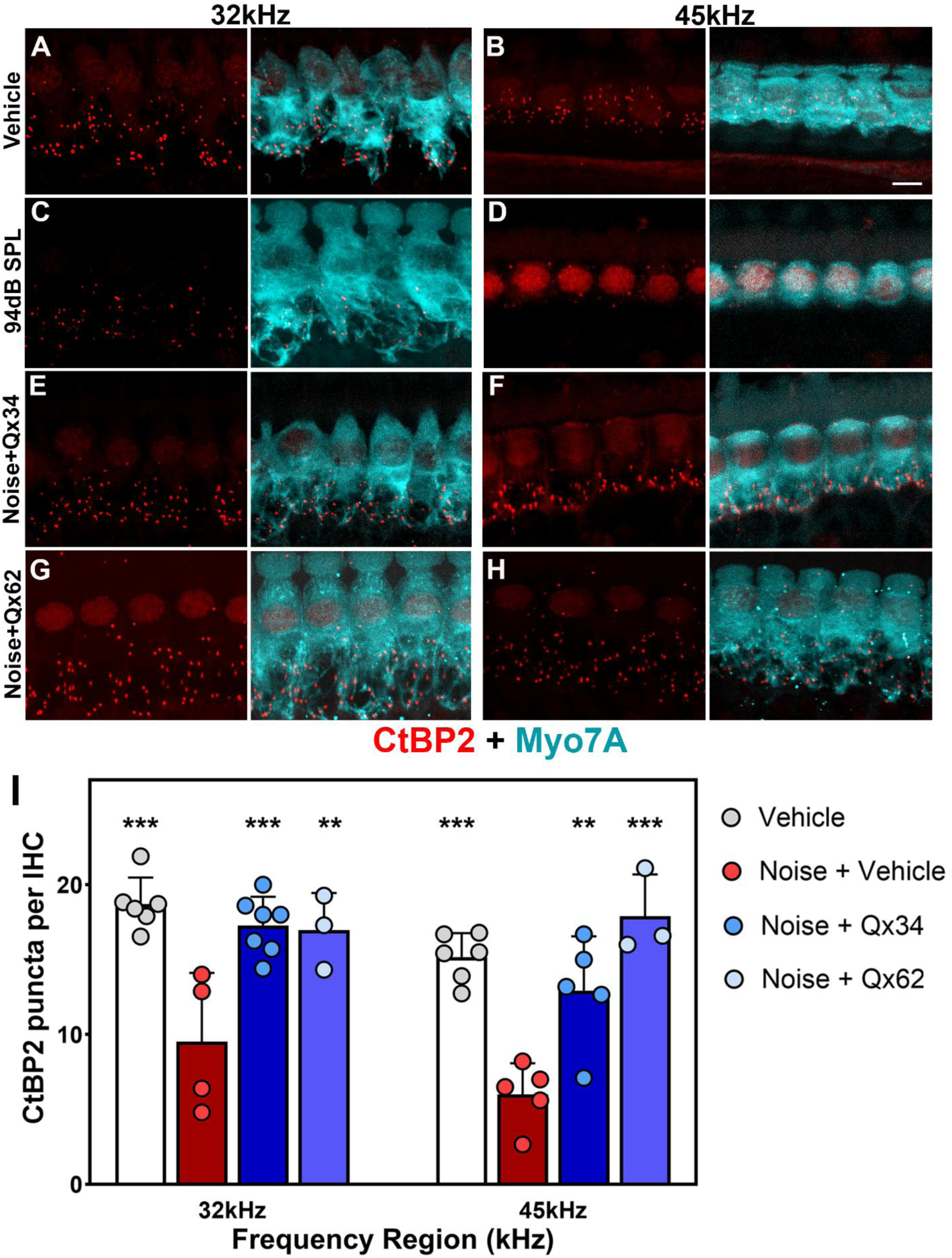
Qx34 and Qx62 protect ribbon synapses after noise-induced synaptopathy. **A-H:** Representative micrographs of IHCs from the 32 kHz (**A, C, E, G**) and 45 kHz (**B, D, F, H**) cochlear regions immunostained for the pre-synaptic marker CtBP2 (red) and the hair cell marker Myo7a (cyan). Scale bar=5 µm. **I:** Quantification of IHC ribbon synapses, expressed as mean±SD. Circles represent individual animals (biological replicates). Statistical analysis was performed using one-way ANOVA followed by Dunnett’s multiple comparisons test. **P<0.01, ***P<0.001 *versus* the Noise+vehicle group.

Overall, of the five Qx derivatives advanced from the zebrafish screen, two - Qx34 and Qx62 - demonstrated significant protection against noise-induced synaptopathy in mice, accelerating recovery of auditory function while preserving synaptic integrity and morphology. These two derivatives were therefore progressed to the final stages of characterization as candidate therapies for NIHL.

### Qx62 protects against noise-induced hair cell death

We next tested the therapeutic potential of Qx34 and Qx62 under noise conditions that cause hair cell death^3 38–40^. Using a similar treatment protocol (**Supp. Figure 3B**), animals were exposed to 112dB SPL at 8-16kHz for 2 hours. A single dose of Qx34 or Qx62 (25mg/kg b.w.) was given immediately after noise exposure and ABRs and DPOAEs were tested before and at 1- and 21-days post-noise. A single dose of Qx34 immediately after noise did not improve hearing (**Fig. 6A-B, E**), regardless of whether outer hair cells (OHCs) were still present (32kHz, **Fig. 6J, N**) or not (45kHz, **Fig. 6K, N**). For Qx62, we observed significant recovery of ABR thresholds only at 45kHz compared with the noise+vehicle group, although a trend toward lower thresholds was evident at the remaining frequencies (**Fig. 6C, D**). Importantly, DPOAEs were significantly lower in Qx62-treated animals compared to those exposed to noise alone (**Fig. 6E**), and we did not observe any significant OHC loss (**Fig. 6K-M**), suggesting a window of opportunity for therapeutic improvement.

**Figure 6.**
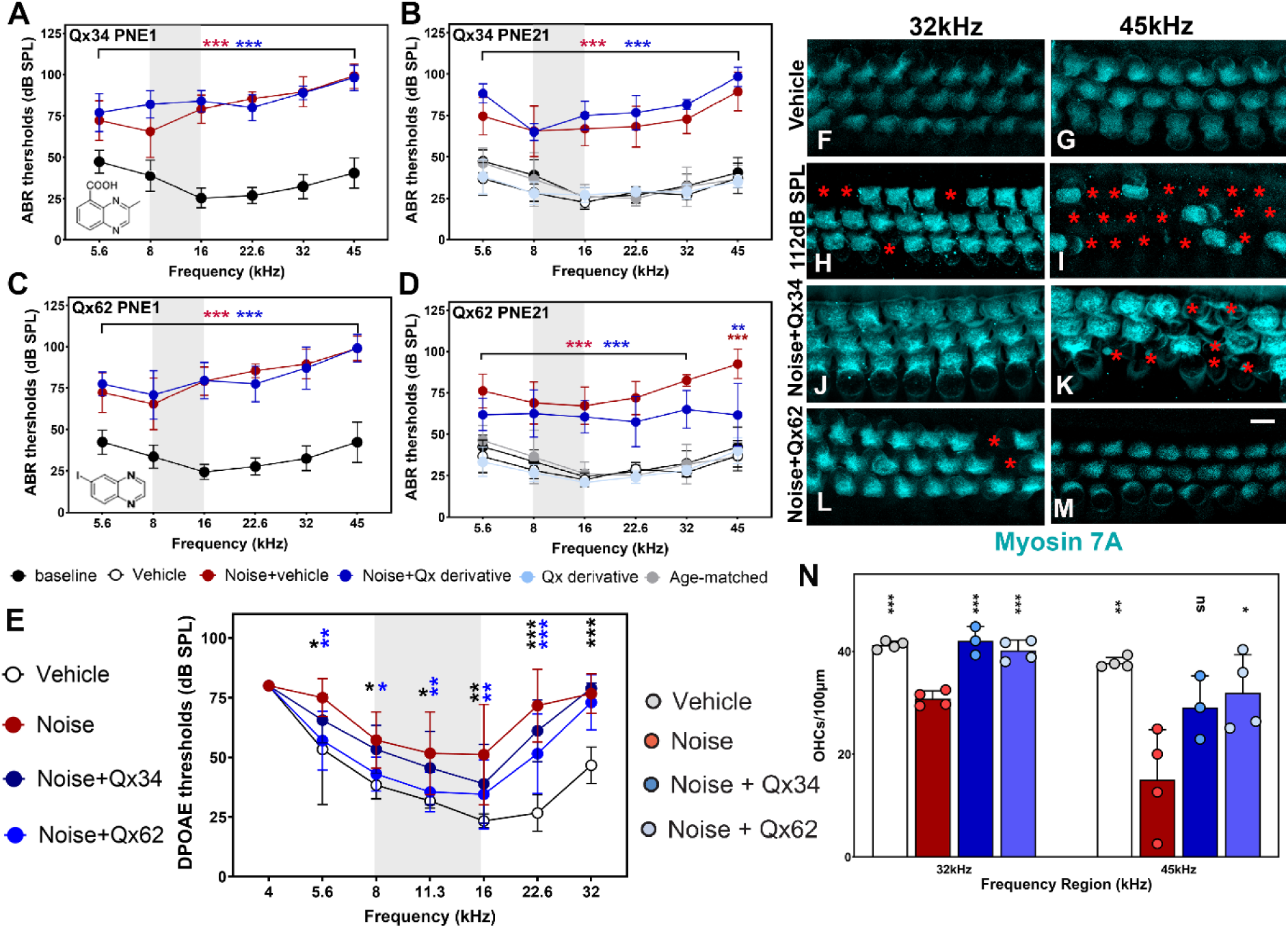
Only Qx62 shows therapeutic potential against noise-induced hair cell loss. Seven- to eight-week-old CBA/CaJ mice were exposed to 112dB SPL octave band noise (8-16 kHz) for 2 hours and immediately received an intraperitoneal injection of corn oil, Qx34, or Qx62 at 25mg/kg b.w. ABR thresholds were measured at 1 day (PNE1) and 21 days (PNE21) after noise exposure. **A-B:** Qx34. **C-D:** Qx62. Results are expressed as mean±SD. Statistical analysis was performed using two-way ANOVA followed by Dunnett’s multiple comparisons test. **P<0.01, and ***P<0.001 for Noise+vehicle (red asterisks) and Noise+Qx derivative (blue asterisks) compared with the baseline/vehicle group. **E:** DPOAE thresholds were measured at PNE21 in vehicle (white dots), Noise+vehicle (red dots), Noise+Qx34 (dark blue dots), and Noise+Qx62 (blue dots) groups. Results are expressed as mean±SD. Statistical analysis was performed using two-way ANOVA followed by Dunnett’s multiple comparisons test. *P<0.05, **P<0.01, and ***P<0.001 for Noise+Qx34 (dark blue asterisks) and Noise+Qx62 (blue asterisks) compared with the vehicle group. **F-M:** Representative micrographs of OHCs from the 32 kHz (**F, H, J, L**) and 45 kHz (**G, I, K, M**) cochlear regions immunostained for the hair cell marker Myo7a (cyan). Scale bar=5 µm. Red asterisks indicate missing OHCs. **N:** Quantification of OHC survival, expressed as mean±SD. Dots represent individual animals (biological replicates). Statistical analysis was performed using one-way ANOVA followed by Dunnett’s multiple comparisons test. *P<0.05, **P<0.01, ***P<0.001 *versus* the Noise+vehicle group. ns = not significant. Group sizes were as follows: Noise+vehicle, N=10; Noise+Qx34, N=8; Noise+Qx62, N=8; Vehicle, N=8; Baseline, N=10. Gray shaded bar in **A-E** indicates the octave band noise.

These results demonstrated the potential of Qx62 to protect cochlear function and structure and justified fruther efforts to optimize its administration protocol for more effective protection against NIHL. Because Qx62 had initially been administered as a single 25 mg/kg. b.w. dose immediately after noise exposure, we next evaluated a repeated-dosing regimen in which Qx62 was given at 50 mg/kg. b.w. once daily for seven consecutive days, beginning immediately after noise exposure (**Supp. Fig. 3C**). Under this regimen, protection improved, reaching statistical significance at all frequencies compared with animals that were also exposed to noise but received corn oil instead (**Fig. 7**). Both ABR thresholds (**Fig. 7A**) and DPOAEs (**Fig. 7B**) were significantly lower than noise-exposed vehicle controls, and cochlear morphology was preserved (**Fig. 7C-H**). While OHC numbers were significantly reduced in the noise-exposed animals at 32kHz and 45kHz frequency regions compared with Vehicle-only group, no significant differences were observed between the Vehicle-only and Qx62-treated groups. These findings indicate that the higher-dose, repeated-dosing protocol was sufficient to protect the inner ear from intense noise damage.

**Figure 7.**
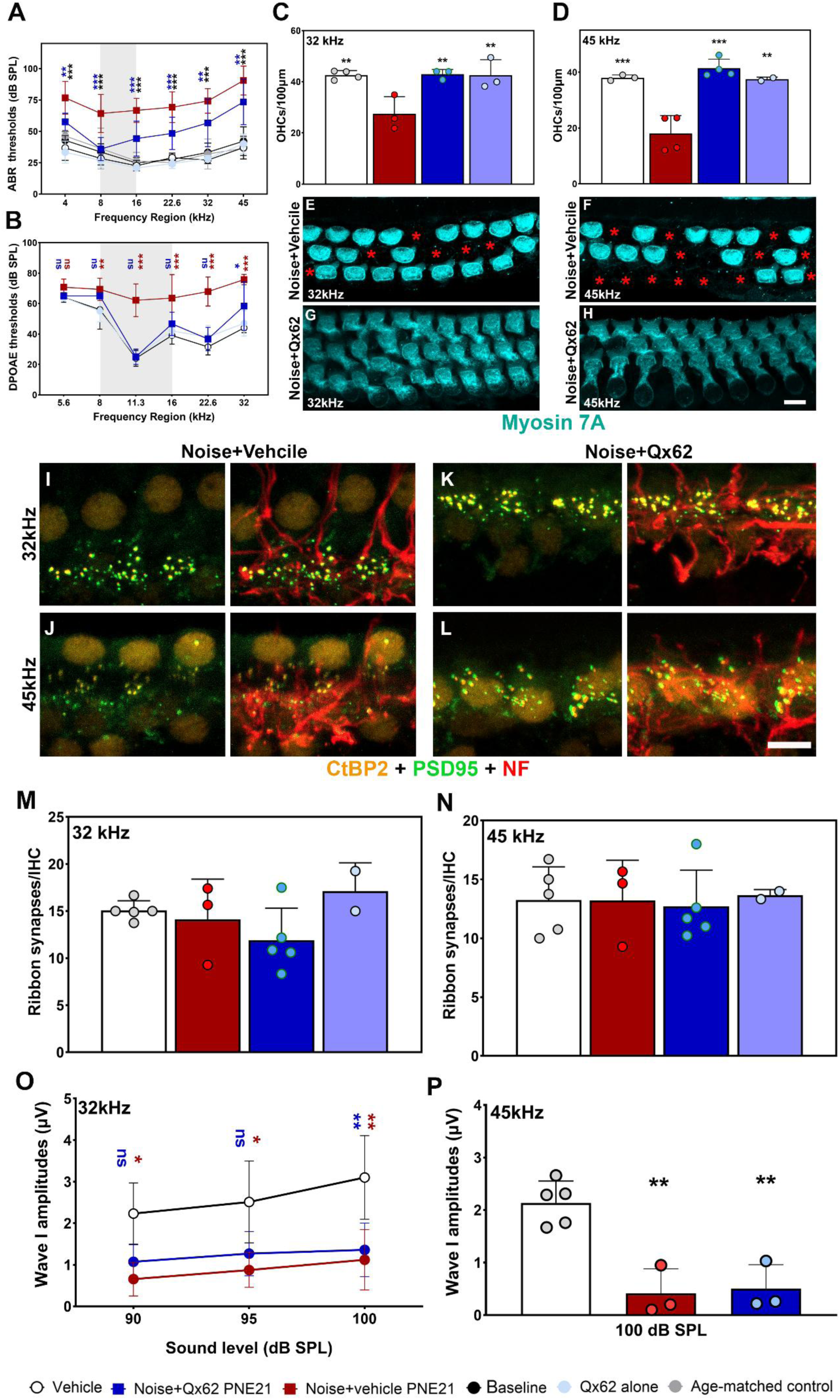
Multidose Qx62 regimen protects against noise-induced cochlear damage. Seven- to eight-week-old CBA/CaJ mice were exposed to 112dB SPL octave band noise (8-16 kHz) for 2 hours and immediately received an intraperitoneal injection of corn oil or Qx62 (50mg/kg b.w.), followed by once-daily injections for an additional six consecutive days. ABR (**A**) and DPOAE (**B**) thresholds were measured 21 days after noise exposure (PNE21). Results are expressed as mean±SD. Statistical analysis was performed using two-way ANOVA followed by Dunnett’s multiple comparisons test. **A:** *P<0.05, **P<0.01, and ***P<0.001 for the Qx62-only group (black asterisks) and the Noise+Qx62 group (blue asterisks) compared with the Noise+Vehicle group. Group sizes were as follows: Noise+vehicle, N=10; Noise+Qx62, N=6; Qx62, N=6. **B:** **P<0.01, and ***P<0.001 for the Noise+vehicle group (red asterisks) and the Noise+Qx62 group (blue asterisks) compared with the Qx62-only/baseline groups. Group sizes were as follows: Baseline, N=10, Noise+Vehicle, N=10; Noise+Qx62, N=5; Qx62, N=5. C-D: OHC counts for the 32 kHz and 45 kHz regions in Vehicle, Noise+Vehicle, Noise+Qx62 and Qx62-only groups. Results are expressed as mean±SD. Statistical analysis was performed using one-way ANOVA followed by Dunnett’s multiple comparisons test. **P<0.01, and ***P<0.001 compared with Noise + Vehicle. **E-H:** Representative images of the organ of Corti at 32 kHz (**E, G**) and 45 kHz (**F, H**) frequency regions for Noise+Vehicle (**E-F**) and Noise+Qx62 (**G-H**) groups. Samples were stained for Myosin 7A (cyan). **I-L:** Representative images of the IHC immunostained for the pre-synaptic marker CtBP2 (orange), the post-synaptic marker PSD95 (green) and the neuronal fiber marker neurofilament (red). **I, K**: 32 kHz region. **J, L:** 45 kHz. **I-J:** Noise + Vehicle group. **K-L:** Noise + Qx62 group. **E-L:** Scale bar= 5µm. **M-N:** Quantification of the ribbon synapses (CtBP2:PSD95 pair) in the 32 kHz and 45 kHz regions. **O-P:** Suprathreshold wave-I amplitudes at 32 kHz (**O**) and 45 kHz (**P**) regions for Noise Vehicle (red), Noise+Qx62 (blue) and Vehicle-only (white) groups. Results are expressed as mean±SD. Statistical analysis was performed using two-way (**O**) or one-way (**P**) ANOVA followed by Dunnett’s multiple comparisons test. **O:** *P<0.05, **P<0.01 for the Noise+Vehicle group (red asterisks) and the Noise+Qx62 group (blue asterisks) compared with the Vehicle-only group. P: **P<0.01 compared to Vehicle-only group. Group sizes were as follows: Noise+vehicle, N=3; Noise+Qx62, N=3; Vehicle-only, N=5. ns= not significant. Dots represent individual animals (biological replicates). Gray shaded bar in **A-B** indicates the octave band noise.

We also assessed IHC ribbon synapse numbers (**Fig. 7I-N**). Comparison of the Noise+Vehicle and Noise+Qx62 groups revealed no significant differences in either CtBP2:PSD95 paired synapses or total synapse numbers. Moreover, neither group differed significantly from Vehicle-only controls. In contrast, analysis of suprathreshold ABR wave-I amplitudes at the 32 kHz and 45 kHz frequency regions showed a significant reduction in the Noise+Vehicle group compared with the Vehicle-only group (**Fig. 7O-P**). Animals exposed to noise and treated with Qx62 also exhibited reduced wave-1 amplitudes; however, this decrease reached statistical significance only at the 45 kHz frequency region. These results are consistent with previous studies demonstrating that, in animal models of permanent threshold shift, reductions in wave-I amplitudes are not longer proportional to synapse counts. In these models, ABR amplitudes decline to a greater extent than would be predicted from synapse survival alone, likely reflecting a combination of neuronal dysfunction and OHC loss^9^. Nonetheless, animals treated with Qx62 performed better than those receiving corn oil alone, suggesting that Qx62 preserves not only cochlear morphology but also neuronal function.

Overall, Qx62 protected against noise-induced hair cell death, providing significant protection across all frequency regions when administered at 50mg/kg b.w for seven consecutive days, whereas Qx34 protected only against ribbon synapse loss caused by lower-intensity noise.

### Qx62 modulates noise-dependent activation of NF-κB pathway in the inner ear

Previous work has shown that Qx possesses anti-inflammatory properties. To assess whether Qx62 shares this property and whether it underlies the protective effects we observed in the inner ear, we ran a 41-plex mouse cytokine/chemokine panel on cochlear samples collected 7 and 21 days post-exposure (**Fig. 8**). We detected 28 metabolites in the mouse cochlea (**Supp. Fig. 5**), of these, 9 differed significantly in abundance between vehicle-treated and Qx62-treated animals exposed to 112dB SPL noise (8-16kHz for 2 hours). Most of these differences were observed 7 days post-noise exposure (**Fig. 8A-B**). IL-4 and IL-7 were significantly increased in noise+Qx62 mice compared to noise+vehicle animals. IL-4 is involved^41^ in the switch from pro-inflammatory M1 to repair-associated M2 macrophages, whereas IL-7 plays a role in lymphocyte homeostasis^7 41–43^. Trophic factors EGF and FGF-2 were also increased, which support cell proliferation and survival^44–46^. Qx62 treatment also significantly reduced IL-17a, IL-22, CXCL1, and CXCL10, cytokines involved in innate immune cell recruitment and the pro-inflammatory response^47–50^. IL-10 was also significantly reduced. Because this cytokine typically rises in proportion to the size of the pro-inflammatory response as a compensatory anti-inflammatory signal, its reduction is consistent with a smaller inflammatory response requiring less counter-regulation^51–53^. Levels of all 9 metabolites returned to baseline levels by PNE21, suggesting that Qx62 suppresses the acute inflammatory response occurring within the first week after acoustic injury (**Fig. 8B**). LIX (CXCL5) and MIG (CXCL9) were also significantly decreased in Qx62-treated animals relative to noise-exposed controls at PNE 21 (**Fig.8C**). Because both cytokines rise during tissue repair, remodeling, and the pro-inflammatory response, this finding is consistent with reduced inflammation and injury following Qx62 treatment.

**Figure 8.**
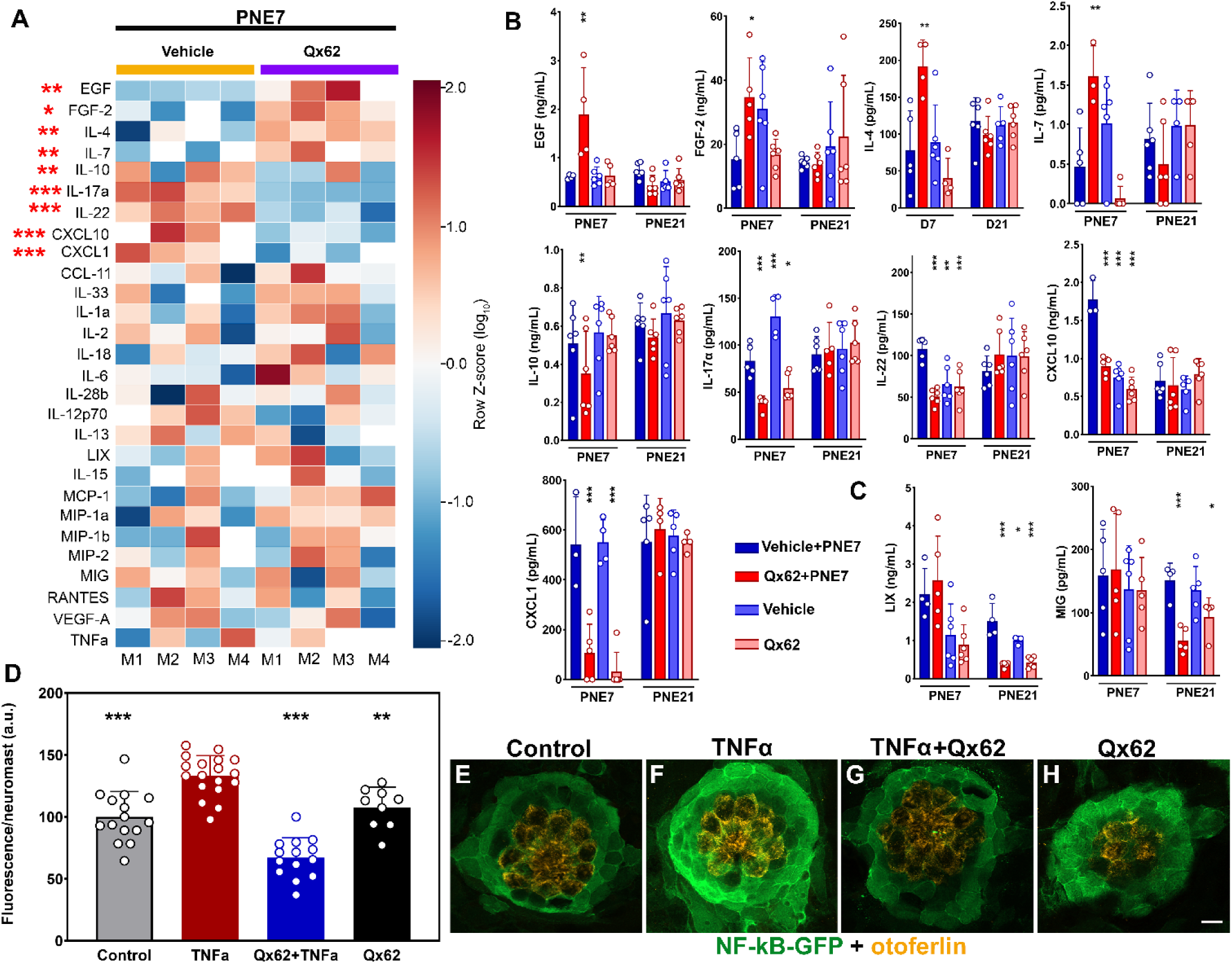
Qx62 protects against noise-induced cochlear damage by attenuating the early inflammatory response. A-C: Seven- to eight-week-old CBA/CaJ mice were exposed to 112dB SPL octave band noise (8-16 kHz) for 2 hours and immediately received an intraperitoneal injection of corn oil or Qx62 (50mg/kg b.w.) followed by once-daily injections for an additional six days. Cochlear tissue was collected at PNE7 and PNE21. **A:** Heatmap of the 28 cytokines/chemokines detected in mouse cochleae using the MILLIPLEX Mouse Cytokine Expansion Panel 1, expressed as log_10_ transformed Z-scores. Asterisks indicate significant differences in the metabolite expression between Noise+vehicle and Noise+Qx62 groups. N=4 animals per group. **B-C:** Absolute abundances of cytokines/chemokines showing significant differences at PNE7 (**B**) or PNE21 (**C**). Each dot represents one animal (biological replicate). Statistical analysis was performed using one-way ANOVA followed by Dunnett’s multiple comparisons test. *P<0.05, **P<0.01, and ***P<0.001 *versus* the Vehicle+PNE7 group. **D:** Quantification of the total fluorescence intensity per neuromast in 5-6 dpf *Tg*(*NFKB:EGFP)* larvae incubated with 10ng/mL of TNFα for 30 minutes or E3 medium, followed by treatment with vehicle (DMSO) or 1nM of Qx62 for 2 hours. Results are expressed as mean±SD. Statistical analysis was performed using one-way ANOVA followed by Dunnett’s multiple comparisons test. **P<0.01, and ***P<0.001 *versus* TNFα-treated group. Each dot represents one zebrafish (biological replicate). **E-H:** Representative micrographs of *Tg*(*NFKB:EGFP)* larvae immunostained for GFP (green) and the hair cell marker otoferlin (orange). Scale bar = 5 µm.

Since most of these differentially expressed cytokines are directly or indirectly associated with the NF-κB pathway^22 23 26 44^, and given that prior evidence suggests Qx directly targets this pathway, we next examined the effect of Qx62 on NF-κB activity using a reporter zebrafish line expressing green fluorescent protein (GFP) under a promoter containing a consensus NF-κB binding sequence (**Fig. 8E-F**). Transgenic *Tg*(*NFKB:EGFP)* zebrafish^19^ were incubated with TNFα (10ng/mL, 30 min) to activate the pathway, followed by a 2-hour incubation in E3 water containing vehicle (DMSO) or Qx62 (1nM). TNFα incubation significantly increased GFP expression, primarily in neuromast supporting cells, indicating NF-κB activation; incubation with Qx62 significantly reduced GFP levels back to control values.

Together, these results suggest that Qx62 protects the inner ear from noise damage by reducing the early inflammatory response and increasing trophic factors that support tissue repair, and that it does so by inhibiting the NF-κB pro-inflammatory response.

## DISCUSSION

No pharmacological agent is currently approved to prevent or treat NIHL, and most experimental strategies act downstream after ROS have already damaged hair cells and synapses^40 54–56^. Our zebrafish-to-mouse discovery pipeline demonstrates that the quinoxaline derivative Qx62 intervenes earlier, protecting the cochlea from both synaptopathic and HC-damaging noise, in association with suppression of the early NF-κB response. Because cochlear NF-κB activation peaks within hours of acoustic trauma and precedes the oxidative and apoptotic phases of injury^24^, a compound capable of attenuating this signal before it propagates is well positioned to preserve cochlear structures that would otherwise be irreversibly lost.

Importantly, NIHL is not a single clinical entity. Moderate, chronic noise exposure primarily damages afferent ribbon synapses while sparing HCs, whereas intense noise exposure also causes OHC death^9^ ^40^. Our two mouse paradigms were designed to model these distinct injury profiles, both of which are clinically relevant. Synaptopathy without HC loss is common and is thought to underline much of the speech-in-noise difficulty reported by individuals with normal audiograms^8 17^, whereas OHC loss contributes to the more severe, threshold-elevating hearing loss typically associated with blast and impulse noise exposure in military settings^1–3 40^. Within this framework, our findings support therapeutic potential for both Qx34 and Qx62. In the synaptopathy model, both compounds accelerate ABR recovery and preserve wave-I amplitude, latency, and presynaptic ribbon counts to a similar extent, suggesting that either compound could mitigate synaptopathy-driven hearing deficits. In contrast, in the higher-intensity noise paradigm specifically designed to induce OHC death, the compounds diverged. Once its dosing regimen was optimized, Qx62 fully protected OHCs and synapse function, whereas Qx34 did not. This distinction is important for the future clinical position of each compound but does not diminish the therapeutic value of Qx34.

Our findings are consistent with a growing body of evidence implicating NF-κB as a central mediator of NIHL pathogenesis. Noise exposure activates the TLR4/NF-κB axis^22 25 26 57^, promoting macrophage recruitment and cochlear inflammation^7 42 58 59^. Independent studies of Qx and chemically related quinazoline scaffolds have identified analogs that inhibit IKKβ and other upstream regulators of NF-κB while exhibiting favorable oral bioavailability and low cytotoxicity^21 60^. These observations raise the possibility that Qx62 acts as a direct inhibitor of NF-κB activation rather than simply scavenging downstream ROS. The reduction of TNFα-induced GFP expression observed in the *Tg(NFKB:EGFP)* zebrafish line following Qx62 treatment supports this interpretation and, together with the mammalian cytokine data, suggests that this mechanism is conserved across both experimental systems used in this study.

The cochlear cytokine profile after Qx62 treatment is consistent with established injury-repair trajectories. Increased IL-4 and IL-7 expression is compatible with a shift from pro-inflammatory M1 macrophages toward a reparative M2-like phenotype^43 61 62^, a transition that promotes resolution of inflammation and tissue remodeling. The concurrent increase in EGF and FGF-2 aligns with previous reports demonstrating that these growth factors support cochlear sensory cell survival, including through NF-κB-dependent pro-survival signaling that is mechanistically distinct from the pathological NF-κB activation associated with acoustic injury^44 53^. This distinction may be important, as it suggests that Qx62 does not globally suppress NF-κB activity but instead selectively dampens maladaptive innate immune signaling while preserving pathways that promote tissue survival and repair. The accompanying reductions in IL-17a, IL-22, CXCL1, and CXCL10 are likewise consistent with suppression of innate immune cell recruitment; IL-17a and CXCL10 are well-established mediators of pro-inflammatory leukocyte trafficking^48 63–65^, whereas IL-22 has context-dependent functions that range from promoting tissue injury to facilitating tissue repair depending on the timing and the inflammatory milieu^48 66^. This duality may explain why reduced IL-22 expression, rather than increased expression, was associated with favorable outcomes in our acute, 7-day post-noise model. The parallel reduction in IL-10, which is typically induced as a homeostatic brake on inflammation, is most parsimoniously explained as a secondary consequence of a diminished inflammatory response requiring less counter-regulation, rather than a loss of anti-inflammatory signaling^52 67 68^.

Because these mechanistic studies were performed only with Qx62, they do not establish whether Qx34 acts through the same pathway. Determining whether Qx34 similarly modulates NF-κB signaling and the cochlear cytokine response will be an important direction for future studies. Since Qx34 was evaluated using only a single post-noise dosing regimen in the OHC-damaging noise paradigm, it remains unresolved whether it is intrinsically less effective at preventing OHC death or, like Qx62, simply requires optimization of its dosing strategy. Addressing this question will be an important priority for future work. Nevertheless, Qx34 consistently protected against synaptopathy, a therapeutically meaningful outcome given the prevalence and functional impact of cochlear synaptic loss independent of HC degeneration. We therefore view Qx34 and Qx62 as complementary therapeutic leads rather than primary and secondary candidates: Qx62 as a candidate for the broader spectrum of noise injury, including severe HC-damaging exposures such as military blast trauma, and Qx34 as a candidate for the synaptopathy associated with more moderate or chronic noise exposure, pending further optimization and evaluation in higher-intensity injury paradigms.

Together, these findings establish a translational discovery pipeline that successfully identified quinoxaline derivatives with robust otoprotective activity across complementary vertebrate models of NIHL. By combining phenotypic screening in zebrafish with mechanistic and functional validation in mice, we identified compounds that preserve cochlear structure and function while modulating early inflammatory signaling upstream of irreversible tissue damage. Although additional studies are needed to define pharmacokinetics, therapeutic windows, long-term efficacy, and molecular targets, our results position Qx34 and Qx62 as promising candidates for pharmacological prevention of NIHL. More broadly, these findings support early modulation of NF-κB-dependent inflammatory signaling as a viable therapeutic strategy for preserving hearing after acoustic trauma.

Although our findings establish Qx34 and Qx62 as promising otoprotective candidates, several limitations should be acknowledged. First, hearing function and cochlear morphology were evaluated only through 21 days after noise exposure. Consequently, we cannot determine whether the protection conferred by either compound is permanent or whether delayed cochlear degeneration may still occur. Long-term studies will therefore be necessary to establish the durability of the observed structural and functional protection. Second, our work focused exclusively on a therapeutic paradigm in which treatment was initiated after acoustic injury. Although this approach reflects the clinical need to treat unexpected noise exposure, it does not address the potential prophylactic efficacy of these compounds. Because many forms of hazardous noise exposure, including occupational, military, and recreational settings, are predictable, an equally important question is whether quinoxaline derivatives administered before noise exposure can prevent cochlear injury altogether. Addressing these questions will provide a more complete understanding of the clinical utility of these compounds across the full spectrum of noise-exposure scenarios.

## MATERIALS AND METHODS

### Study Design

The aim of this study was to identify Qx derivatives as potential therapeutics for NIHL. Sample sizes were determined on the basis of previous hearing studies conducted in our laboratory ^6 36^. Following baseline auditory testing, animals were allowed to recover from anesthesia for 2-3 days before noise exposure and drug treatment. Mice were randomly assigned to experimental groups with approximately equal numbers of females and males. Inner ears were collected 7 or 21 days after noise exposure for morphological analysis or Luminex multiplex cytokine assay.

Individual zebrafish or mice were treated as independent biological replicates.

### Animals

*Z*ebrafish and mice were maintained at the Creighton University Animal Resource Facilities in accordance with Institutional Animal Care and Use Committee (IACUC) guidelines (IACUC protocols 1192 and 1123).

For drug screening, 5-6 dpf wild-type TuAB zebrafish larvae were used. Pre-synaptic ribbon and afferent neuron studies we performed using 5-6 dpf *Tg(NeuroD-GFP)*^37^ larvae. In vivo studies of NF-κB signaling were performed using the *Tg*(*NFKB:EGFP)*^18^ reporter line.

Zebrafish were maintained at 28.5°C in E3 media (5mM NaCl, 0.17mM KCl, 0.33mM CaCl2 and 0.33 mM MgSO4, pH 7.2) under a 14-hour light/10-hour dark cycle.

Male and female CBA/CaJ mice (7-8 weeks of age) were used for all noise exposure studies. Mice were housed under a 12-hour light/12-hour dark cycle with *ad libitum* access to food and water. Food supplements and hydrogel were placed in the cages one week before noise exposure.

Every effort was made to minimize animal suffering and reduce the number of animals used.

### Quinoxaline derivatives preparation

For zebrafish experiments, quinoxaline derivatives (**Supp. Fig. 1**) were previously synthesized in-house^19^. Stock solutions were prepared in DMSO at 1,000X concentration and stored at -20°C.

For mouse studies, Qx3, Qx17, Qx23, Qx34 and Qx62 were purchased from Ambeed, Inc. and suspended in corn oil (Sigma-Aldrich, C8267). Suspensions were prepared freshly, sonicated at 37°C for 20 minutes, and mixed thoroughly immediately before intraperitoneal (IP) injection.

### Zebrafish experiments

#### Drug screening^5^

Five- to six-dpf TuAB larvae were exposed to 600 µM kainic acid for 60 minutes at 28°C, followed by a 2-hour incubation with individual quinoxaline derivatives (1nM to 100 µM). After 30–60-minute recovery in E3 medium, larvae were fixed, immunostained for otoferlin (DSHB, HSC-1) and processed for fluorescence microscopy as previously described. Control animals were incubated with DMSO (vehicle) alone. HCs were manually quantified in neuromasts MI1, MI2 and O2 using a Zeiss AxioSkop 2 fluorescence microscope equipped with a 40x oil-immersion objective.

#### Presynaptic ribbon assessment^5^

Five- to six-dpf *Tg(NeuroD-GFP)* larvae were incubated with 600 µM kainic acid for 60 minutes, followed by a 30-minute incubation with the corresponding Qx derivative at the lowest concentration that showed protection in the drug-screening assay. Images were acquired using a Zeiss LSM 700 confocal microscope with a 63X oil-immersion objective. Presynaptic puncta were manually quantified and expressed as number of pre-synaptic ribbons per neuromast.

#### Qx62 studies in the NF-κB reporter line^19^

Five- to six-dpf *Tg*(*NFKB:EGFP)* larvae were incubated with 10 ng/mL of TNF-α (Cell Signaling Technology, 5178) for 30 minutes to activate NF-κB pathway, followed by a 2-hour incubation with Qx62 (1 nM) or DMSO (vehicle) to permit GFP expression. Larvae were then fixed, prepared for confocal microscopy, and GFP fluorescence per neuromast was quantified using ImageJ as previously described.

### Mouse experiments

#### Quinoxaline derivative efficacy studies

Following baseline hearing assessments (ABR and DPOAE), 7-8 week-old CBA/CaJ mice were randomly assigned to one of five groups: untreated controls, vehicle-treated controls (single IP injection of corn oil without noise exposure), noise-exposed vehicle-treated animals, quinoxaline derivative-treated noise-exposed animals (single IP dose), and quinoxaline derivative-treated non-noise controls. Quinoxaline derivatives were freshly prepared in corn oil and administered immediately after noise exposure at 25 mg/kg b.w. The dose was selected on the basis of preliminary studies using the parent quinoxaline compound.

#### Seven-day Qx62 treatment

Following baseline hearing assessment, mice were exposed to noise (112dB SPL, 2hrs, 8-16kHz) and immediately received Qx62 (50 mg/kg b.w., IP). Treatment was repeated once daily for six additional consecutive days. Fresh Qx62 suspensions were prepared each day.

#### Noise exposure

CBA/CaJ mice were exposed to noise as previously described. Briefly, awake animals were placed individually in open-walled containers and exposed to either 94dB SPL (synaptopathy model) or 112dB SPL (permanent hearing loss model) noise for 2 hours (8-16 kHz). Noise was delivered through an exponential horn fitted to a titanium horn driver (JBL 2426H) powered by a Crown XTi1000 amplifier using an RPvdsEx circuit and a RZ6 Multi-I/O processor (Tucker-Davis Technologies - TDT). Before each exposure session, sound levels were calibrated using a Brüel & Kjær sound level meter type 2270 with a microphone type 4044B (No. 3312994) that was pre-calibrated with a sound calibrator (Brüel & Kjær DP0775). Noise exposures were performed between 11:00 AM and 1:00 PM to minimize circadian variation in noise susceptibility. Exposure paradigms were based on previously published studies.

#### Hearing assessment

For the synaptopathy model, ABRs and DPOAEs were recorded using a TDT system as previously described. Measurements were obtained before noise exposure and at 1, 7 and 21 days afterward. Mice were then euthanized, and the inner ears were microdissected for immunohistochemistry studies.

For the OHC injury model, ABRs and DPOAEs were recorded using a National Instruments (NI) system. Hearing assessments were performed before noise exposure and at 1 and 21 days post-noise exposure, followed by tissue collection.

### Luminex assay

Animals were randomly assigned to four groups consisting of noise-exposed (112dB SPL, 2 hours, 8-16 kHz) or no noise controls receiving either corn oil or Qx62 (50mg/kg b.w.) for 7 days. The inner ears were microdissected under cold conditions at 7 and 21 days after noise exposure and cochlea lysates prepared in RIPA buffer (Tris:HCl 50 mM, NaCl 150 mM, NP-40 1%, Na-deoxycholate 0.5%, SDS 1%, EDTA 1 mM, pH 7.5) supplemented with protease and phosphatase inhibitors. Cytokines were quantified using a MILLIPLEX Mouse Cytokine Expansion Panel 1 (MCYT1-190K-SPX, Millipore Sigma) on a LUMINEX L200 analyzer according to the manufacturer’s instructions.

### Immunohistochemistry

#### Zebrafish

Following treatment, larvae were fixed and immunostained using mouse anti-otoferlin (DSHB, HCS-1; RRID:AB_10804296; 1:200), rabbit anti-ribeye B (^69^, RRID: AB_2307328, 1:400), and goat anti-GFP (Novus Biologicals, NB100-1770, RRID:AB_10128178 1:400).

#### Mouse

Cochlear wholemounts were prepared as previously described and immunostained using mouse anti-CtBP2 (BD Biosciences, 612044; RRID:AB_399431; 1:300), alpaca anti-PSD95-FluoTag-X2 (SYSY, N3702-At488-L; RRID: AB_3076105; 1:300), rabbit anti-myosin 7a (Proteus Biosciences, 25-6790; RRID:AB_2314838; 1:400) and chicken anti-neurofilament (Sigma-Aldrich, AB5539, RRID:AB_11212161, 1:1,000).

### Confocal analysis

Zebrafish samples were mounted in ProLong Gold (Thermo Fisher Scientific), and images were acquired using Zeiss LSM 700 or Zeiss LSM 980 confocal microscopes equipped with 63X oil-immersion objectives.

Mouse cochlear whoe mounts were mounted in ProLong Gold and imaged using a Zeiss 980 LSM confocal microscope.

Z-series images were collected using a 63x objective (NA 1.4) and processed with ZEN Black software. Maximum intensity *Z*-projections were used for analysis.

### Quantification

#### ABR and DPOAE analysis

For synaptopathy experiments, thresholds were manually determined as the lowest stimulus intensity at which wave-I was visually identifiable. Wave-I amplitudes were marked as the amplitude (µV) between the highest point of peak one (P1) and the lowest point of the trough (N1). Wave-I latencies were manually marked as the time (ms) from stimulus onset to the P1 peak.

For permanent hearing loss experiments, thresholds, amplitudes, and latencies were determined semi-automatically using the ABR_Peak_Analysis application. ABR and DPOAE thresholds, together with wave-I amplitudes and latencies, were extracted using a custom Python script (https://github.com/ZallocchiLab/ABR-Processor, https://github.com/ZallocchiLab/DP-Processor) *OHCs and ribbon synapse quantification:* Confocal Z-stacks were acquired using 63X oil-immersion objective (NA:1.4), 1.5X digital zoom, and 0.3 μm optical sections extending from the apical surface of the HCs to nerve terminals within the *habenula perforata* beneath the IHCs. OHCs were counted manually and expressed as the number of OHCs per 100 µm. Presynaptic ribbons and postsynaptic densities were also quantified manually. Fluorescence thresholds were adjusted to minimize background while preserving positive signal. Total CtBP2 puncta, total PSD95 puncta, and paired CtBP2/PSD95 puncta were expressed per IHC. Tonotopic frequency mapping was performed with ImageJ as previously described^36^.

#### Fluorescence quantification

GFP fluorescence in *Tg*(*NFKB:EGFP)* larvae was quantified using ImageJ as previously described and expressed as total fluorescence intensity per neuromast.

### Statistical Analysis

Statistical analyses were performed using GraphPad Prism version 10.0.2. Unless otherwise specified, data are presented as mean ± SD. One-way or two-way ANOVA was used, as appropriate, followed by suitable *post hoc* test when significant main effects or interactions were detected. Details regarding sample sizes, experimental replicates, statistical analyses, and error bars are provided in the figure legends. Statistical significance was defined as P≤0.05.

## AUTHOR CONTRIBUTIONS

L.B., I.E., V.M., D.G., A.C. and M.Z. were responsible for investigation. K.F. was responsible for developing the python script. L.B. and M.Z. were responsible for data curation. M.Z. was responsible for conceptualization, analysis performance, development of methodology, and writing of the manuscript.

## ACKNOWLEDGMENTS

We thank the Animal Resource Facility staff, Lyudmila Batalkina, and Linda Goodman for animal husbandry. We also want to thank Drs. Sarath Vijayakumar and Jonathan Fleegel for their help in setting up and calibrating the NI system.

Part of this research was conducted within the AVT Core and Confocal Imaging Facilities at Creighton University, Omaha, NE (RRID:SCR_023866) and supported by a CoBRE Award GM139762 from NIGMS (NIH).

