## Supplementary figures for "Resolving early cochlear inflammation prevents lasting damage from noise exposure"

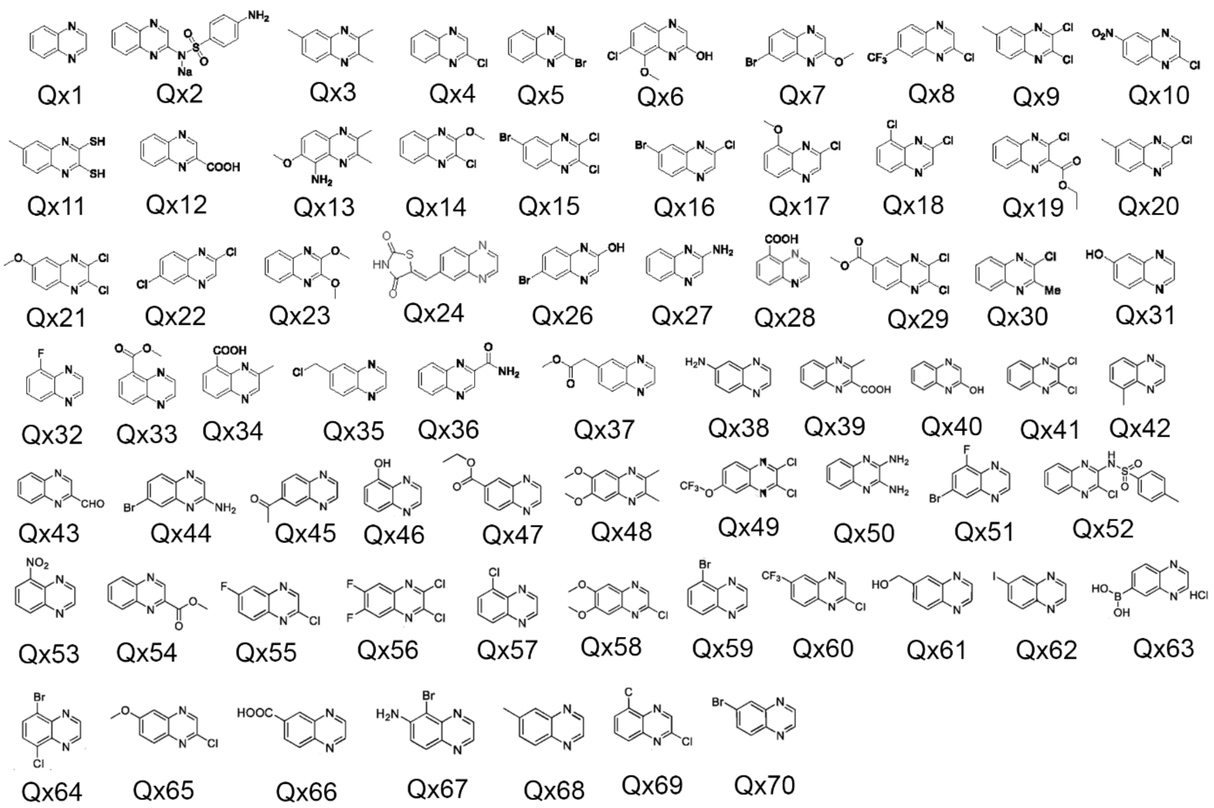


**Figure S1**. *Chemical structure of quinoxaline derivatives*.

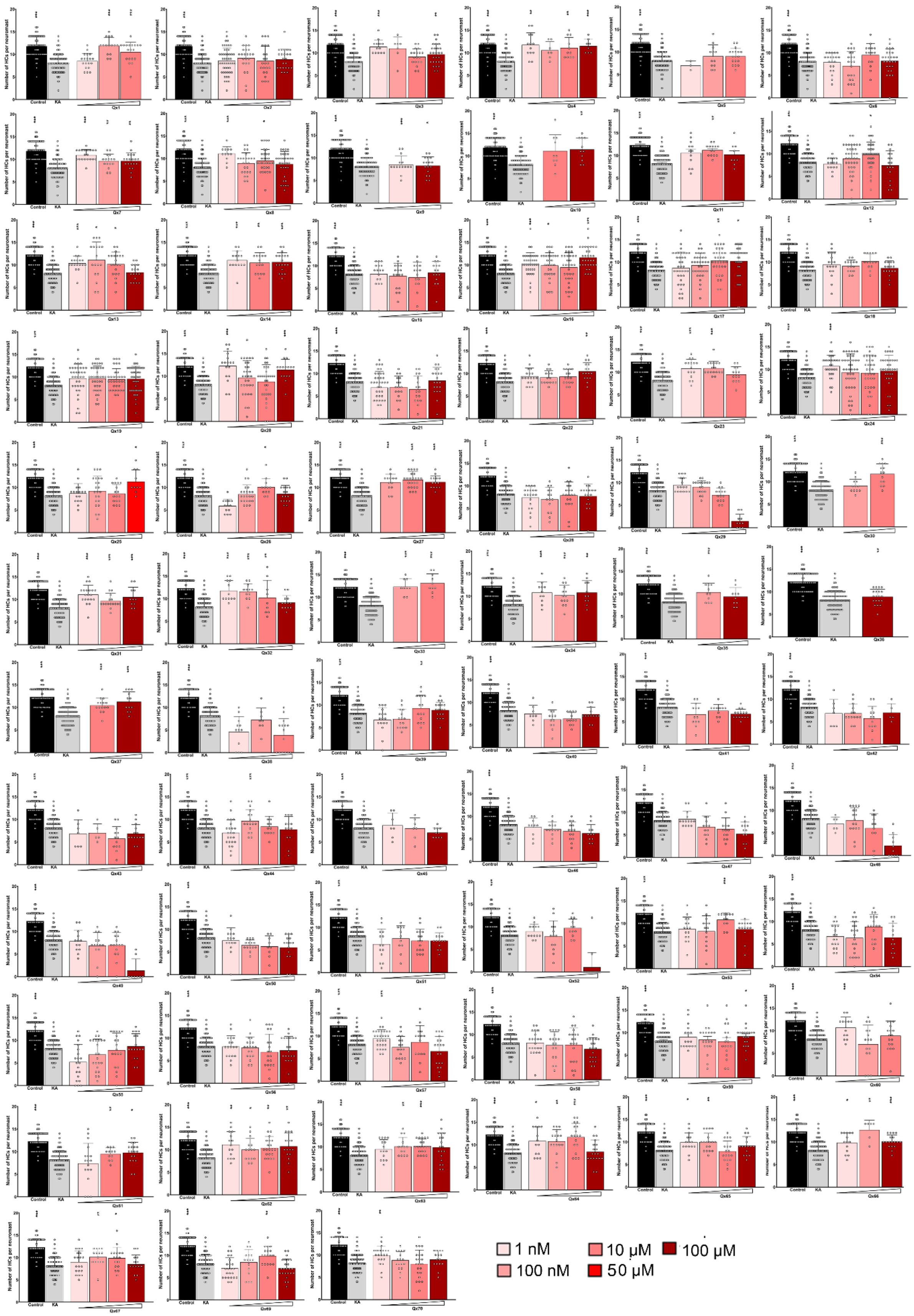


**Figure S2.** *Screening of quinoxaline derivatives in a zebrafish excitotoxic model.* Five- to six-day-post-fertilization zebrafish were incubated with 600 μM kainic acid for 1 hour, followed by a 2-hour incubation with one of the quinoxaline derivatives (Qx1-Qx70) at four concentrations ranging from 1 nM to 100 μM. Neuromast hair cells were immunostained for otoferlin and quantified under a fluorescence microscope. Results are expressed as the number of HCs per neuromast. Each dot represents one neuromast. Black bar indicates vehicle-(DMSO)treated larvae. Grey bar indicates kainic acid only group. Red gradient bars indicate the different concentrations of the Qx derivative used from lowest to highest. Statistical analysis was performed using one-way ANOVA followed by Dunnett's *post hoc* test. N=5-6 fish per group.

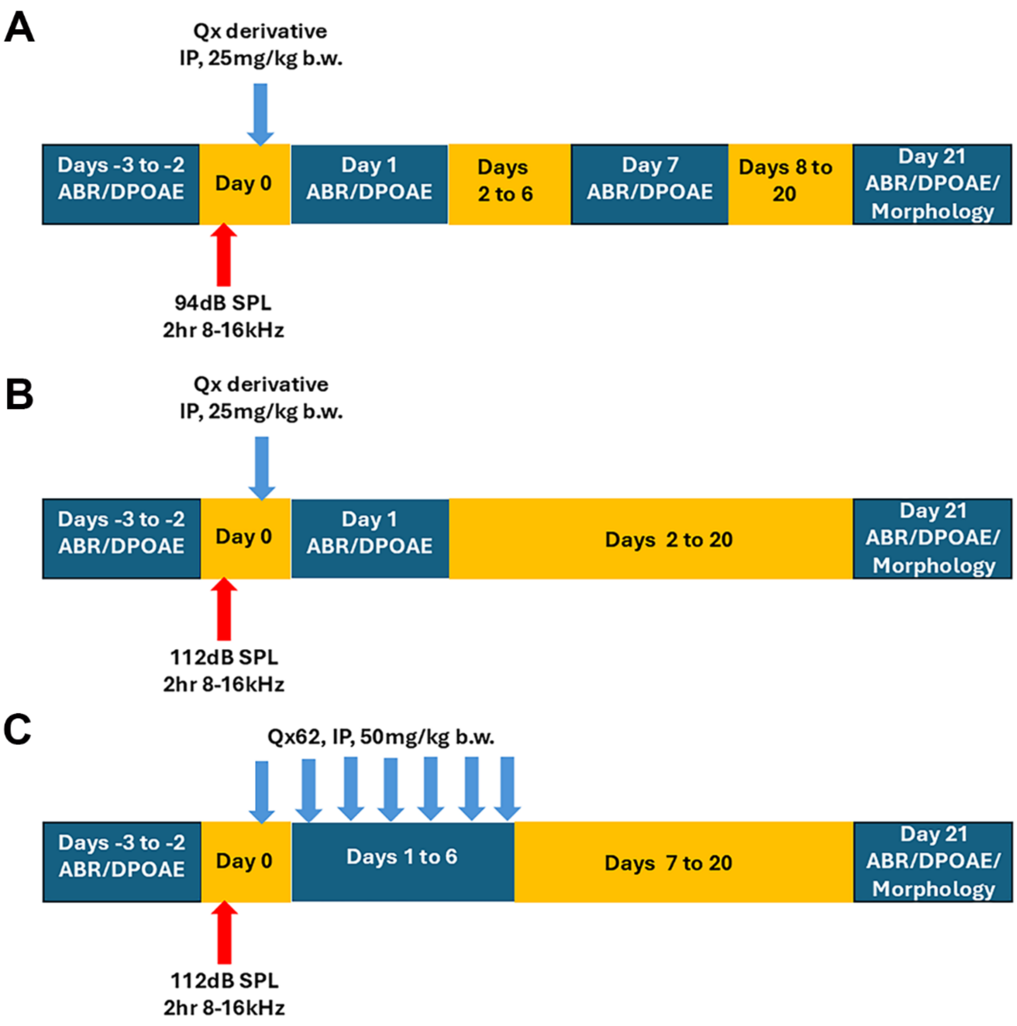


**Figure S3.** *Administration protocols*. **A:** Administration protocol for a synaptopathy model. **B:** Administration protocol for a model of permanent threshold shift elevations due to OHC loss. **C:** Multidose administration protocol for a model of permanent threshold shift elevations.

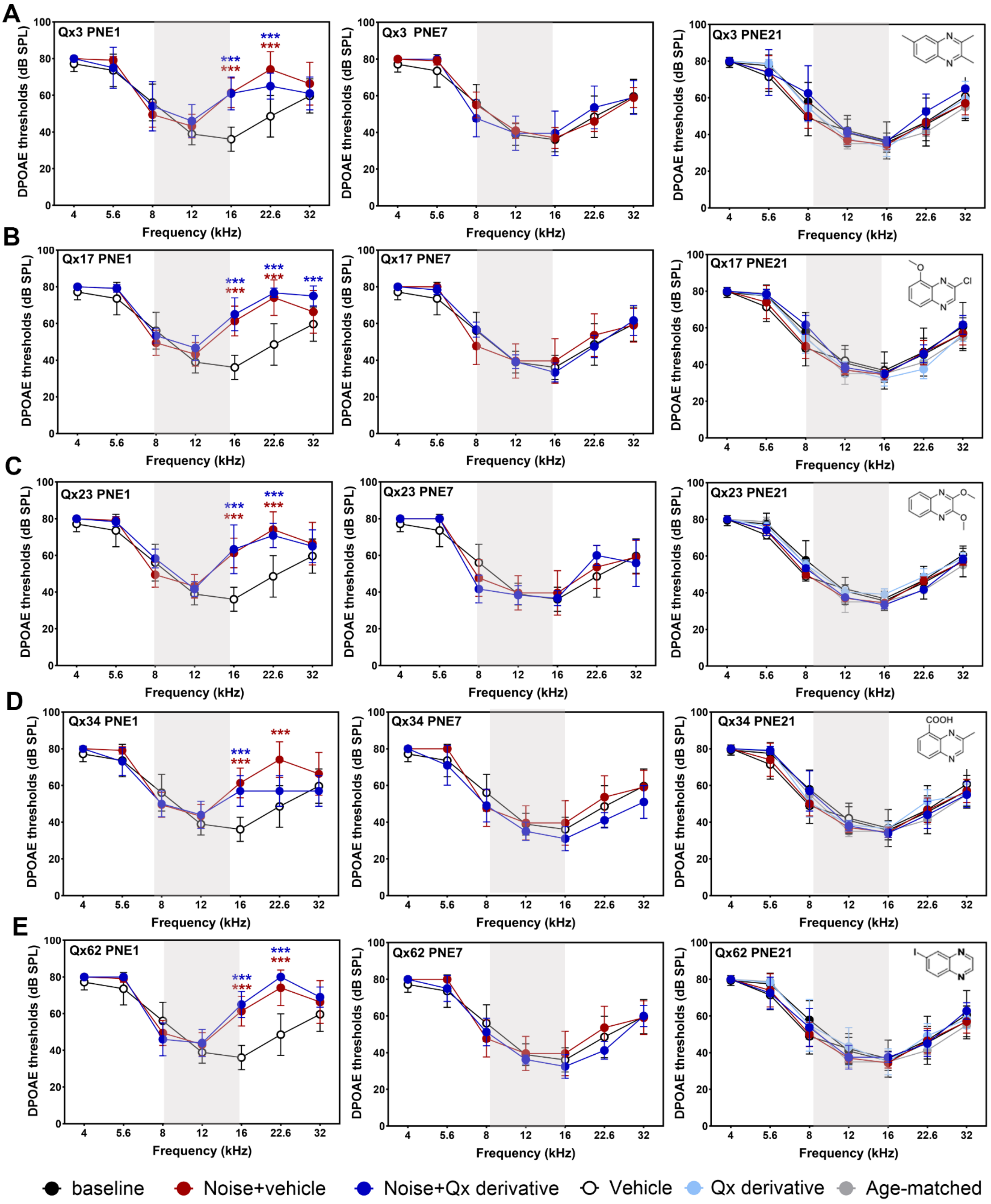
A
Qx3 PNE1
Qx3 PNE7
Qx3 PNE21
B
Qx17 PNE1
Qx17 PNE7
Qx17 PNE21
C
Qx23 PNE1
Qx23 PNE7
Qx23 PNE21
D
Qx34 PNE1
Qx34 PNE7
Qx34 PNE21
E
Qx62 PNE1
Qx62 PNE7
Qx62 PNE21
***
DPOAE thresholds (dB SPL)
0
20
40
60
80
100
Frequency (kHz)
4
5.6
8
12
16
22.6
32
Cl
COOH
I
baseline
Noise+vehicle
Noise+Qx derivative
Vehicle
Qx derivative
Age-matched

**Figure S4.** *DPOAE thresholds recovery after noise-induced synaptopathy*. Seven- to eight-weeks old CBA/CaJ mice were exposed to 94dB SPL for 2 hours at 8-16 kHz and immediately IP injected with corn oil or one of the Qx derivatives at 25mg/kg b.w. DPOAE thresholds were measured at one (PNE1), seven (PNE7), and 21 (PNE21) days post-noise exposure. **A:** Qx3. **B:** Qx17, **C:** Qx23, **D:** Qx34 and **E:** Qx62. Results are expressed as mean±SD. Two-way ANOVA followed by Dunnett's post-test for multiple comparisons. ***$P<0.001$ Noise+vehicle (red asterisks) and Noise+Qx derivative (blue asterisks) *versus* Vehicle/baseline. Number of animals: Noise+vehicle=11. Noise+Qx3=5. Noise+Qx17=6. Noise+Qx23=6. Noise+Qx34=5. Noise+Qx62=6. Vehicle=8. Qx derivative alone=6. Baseline=12. Age-matched=5. Gray shaded bar indicates the octave band noise.

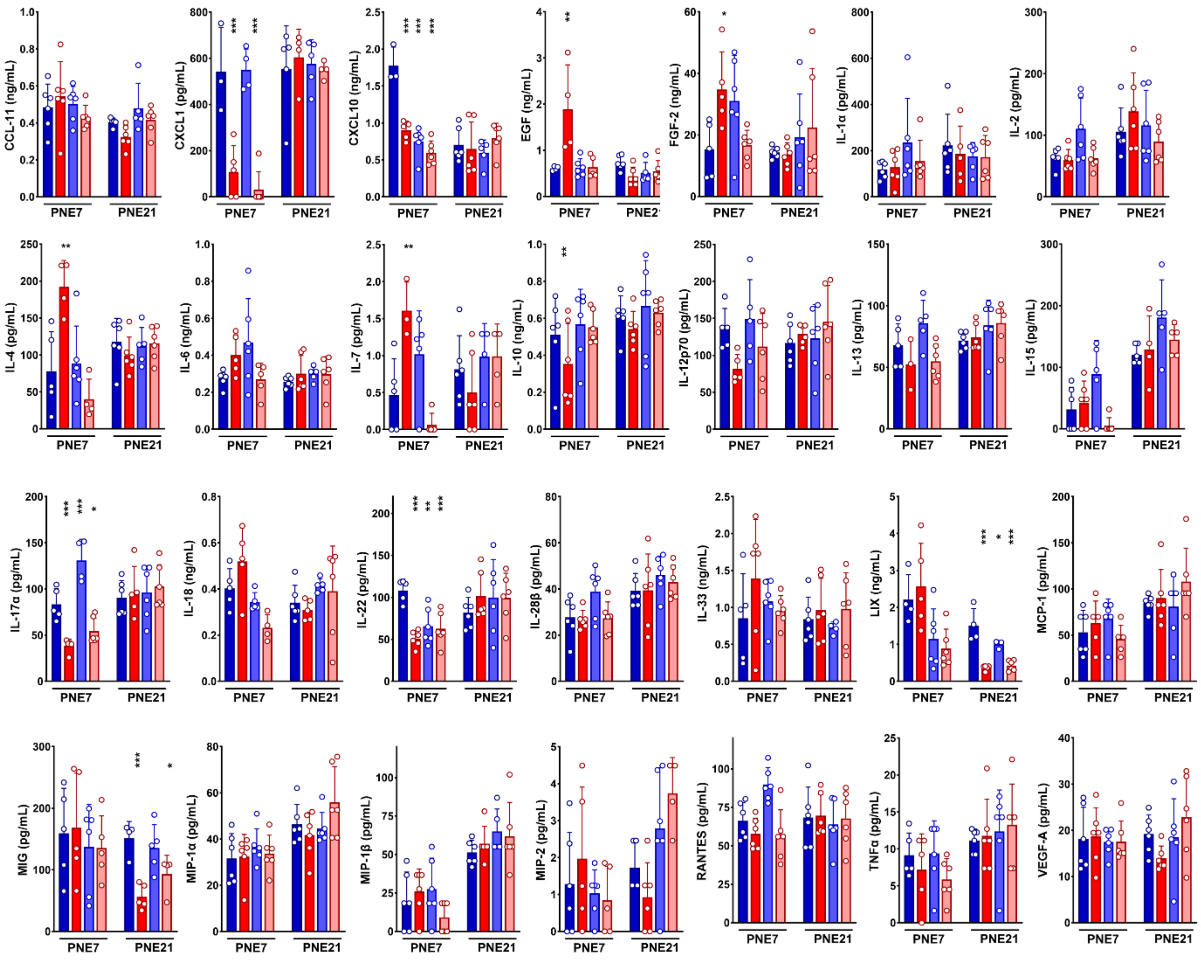


**Figure S5.** *Cytokine/chemokine panel analysis*. Absolute values of the 28 cytokines/chemokines/trophic factors detected in the mouse cochlea for the four groups (Vehicle-only, Noise + Vehicle, Noise + Qx62, and Qx62-only) at 7- and 21-days post-noise exposure (PNE7 and PNE21, respectively).
